# Concepts and Software Package for Efficient Quality Control in Targeted Metabolomics Studies – MeTaQuaC

**DOI:** 10.1101/2020.01.10.901710

**Authors:** Mathias Kuhring, Alina Eisenberger, Vanessa Schmidt, Nicolle Kränkel, David M. Leistner, Jennifer Kirwan, Dieter Beule

## Abstract

Targeted quantitative mass spectrometry metabolite profiling is the workhorse of metabolomics research. Robust and reproducible data is essential for confidence in analytical results and is particularly important with large-scale studies. Commercial kits are now available which use carefully calibrated and validated internal and external standards to provide such reliability. However, they are still subject to processing and technical errors in their use and should be subject to a laboratory’s routine quality assurance and quality control measures to maintain confidence in the results. We discuss important systematic and random measurement errors when using these kits and suggest measures to detect and quantify them. We demonstrate how wider analysis of the entire dataset, alongside standard analyses of quality control samples can be used to identify outliers and quantify systematic trends in order to improve downstream analysis. Finally we present the MeTaQuaC software which implements the above concepts and methods for Biocrates kits and creates a comprehensive quality control report containing rich visualization and informative scores and summary statistics. Preliminary unsupervised multivariate analysis methods are also included to provide rapid insight into study variables and groups. MeTaQuaC is provided as an open source R package under a permissive MIT license and includes detailed user documentation.

---

Targeted mass spectrometry profiling is the workhorse of metabolomics methods. Due to optimization and validation of a defined set of metabolites, it allows for comprehensive routine metabolomics applications such as the analysis of larger cohorts ^1–4^. By focusing on a defined set of metabolites, targeted methods enable both absolute quantification and carry a high potential for standardization by the use of kits validated for specific use cases and sample matrices. However, comprehensive studies require consistent processing and reliable instrumentation to minimize technical variance and interference. Furthermore systematic trends and deviations must be identified and should be quantified ^5–7^.

While standardized methods such as the Targeted Metabolomics Kits of Biocrates ^8^ promise consistent, reproducible and comparable measurements, they are not fully resistant to external influences. These include sample to sample concentration differences, sample handling and processing errors, contamination, sample carryover, batch effects, intra-batch drift, edge effects, missing values of unknown origin and instrument condition ^5,7,9–11^. Consequently, multiple checks and controls are required to verify data quality, consistency and reproducibility (i.e. variance between technical replicates). In particular, comprehensive studies and routine applications benefit from the validation of consistent quality and quantity in and between batches. Standardized targeted kits are well validated and use multiple quality assurance features but vendor software often concentrates on single-analyte or single-sample validation, not harnessing the full quality control potential. For instance, Biocrates MetIDQ software mainly focuses on validation of single analyte accuracy and calibration standard samples (with respect to expected concentration) or single analyte overviews of, for instance, concentration, retention time, peak area (LC) and intensity (FIA). Time-dependent instrument performance monitoring is performed via QC trends and inter-kit measurement per analyte.

Here we briefly outline some potential quality assurance and quality control procedures that can be implemented when using standardized kits for targeted studies. First, we identify various sources of measurement error and then propose methods to detect these errors. Our practical software implementation focuses on Biocrates kits that are widely used ^12–20^. We introduce MeTaQuaC, an easy to use R-package which generates a comprehensive HTML report with interactive elements. The software enables individuals to implement the proposed quality control checks quickly and easily. We envisage that such reports can become part of quality reporting for the community, e.g. as supplementary material in publications.

## QC Measures and MeTaQuaC Software

Quality assurance (QA) and quality control (QC) are important processes to ensure robust data acquisition and to maintain confidence in analysis results. Their importance gains increasing recognition in the metabolomics community ^5,21,22^ and they are required for specific applications by governmental institutions such as the European Medicines Agency (EMA) and the U.S. Food and Drug Administration (FDA) who provide specific guidelines ^23,24^. Put simply, while QA defines processes planned and performed to fulfill defined quality requirements, QC comprises measures to report whether these quality requirements have been met. Here we focus on the later.

Ignoring human error, all experimental uncertainty is due to either random errors or some kind of systematic error including biases and confounders. Random errors are statistical fluctuations (usually in either direction) in the measured data due to the intrinsic precision or other technical limitations of the measurement device or assay. Systematic errors, by contrast, are inaccuracies that occur consistently (usually in the same direction or some form of pattern). Systematic errors may be due to a problem which persists throughout an entire experiment, or which affects only a single batch or subset of samples. Systematic errors can include biases (e.g. due to collection site) in the data collection or trends such as changes in temperature of a laboratory over the course of the day.

The primary aim of quality control measures is to determine that measurements are precise, accurate and reproducible. One necessary consideration is to check for both potential systematic errors as well as unusually large random fluctuation or suspicious measurement ranges. Due to the complexity of the technologies and analyses pipelines, this cannot be achieved by just calculating and thresholding a few key indicators. Thus, we designed a workflow around a comprehensive set of descriptive statistics and visualizations of potential indicators for quality with a specific focus on systematic deviations as well as inspections of calibration and normalization across different runs and batches and data consistency in general.

Spotting systematic error takes a lot of care and effort, especially if the error does not occur in a linear fashion. Systematic error should both be minimized where possible (through instrument maintenance, careful experimental design and laboratory monitoring) and be quantified. Correction of systematic error may be possible in a downstream analysis if reliable quantification was possible. E.g. many batch-correction algorithms exist which use data-driven, internal standards (IS)-based or quality control samples (QC)-based normalization ^5,25–28^. Random errors should be statistically quantified by analysis of suitably defined replicates throughout a study, batch or run. This has been used to exclude variables where the technical variation is greater than the biological signal ^5,27^. However, analysis of technical replicates can be used for additional purposes including estimates of technical variability (between sample, run, batch, etc.), which are an essential input for power calculations given expected effect sizes. For MS based metabolite measurements, the high frequency of missing values pose an additional challenge and must be carefully evaluated in the QC.

Beside biological samples and compounds of interest, a viable targeted method employs several supporting samples as well as supporting analytes as quality assurance measures. Internal standards (IS), i.e. additional compounds, added in known amounts to the samples, allow for IS-based normalization which can assist in correcting for spray intensity or other analysis differences and thus promote comparable analysis independent of instrument or lab. Pooled QC samples created by mixing a number of representative biological samples support the assessment of data quality within and potentially between batches and studies ^5,29^. Reference QC samples consisting of standard reference materials with known concentrations may additionally support inter-study and inter-laboratory analysis ^5^. While reference QC samples provide the benefit of evaluating compound accuracy (with respect to expected concentrations), pooled QC samples have the benefit of matching the same chemical matrix of the samples to be analyzed, thus enabling an assessment of matrix effects. Both enable the evaluation of compound measurement reproducibility (repeatability precision) ^5^ and can be used to correct for systematic errors via QC-based normalization. Calibration standard samples of varying known compound concentrations allow calculation of calibration curves and thus enable absolute quantification of compounds in other samples (with due regard for any differences in matrix effects if sample matrices are different to the EDTA plasma that Biocrates was developed for).

Biocrates adopts a combination of liquid chromatography-mass spectrometry (LC-MS) analysis and flow injection analysis-mass spectrometry (FIA-MS, also known as direct infusion mass spectrometry or DIMS) for their kits. For the LC-MS analysis, Biocrates uses isotopic variants of their target compounds as internal standards with analogous analytical performance which allows for compound-specific normalization within each sample (IS-based normalization). Seven calibration standard samples are available to enable absolute quantification of several compounds (varying with the kit). However, FIA-MS measurements are limited to normalization by internal standards alone (referred to as one-point calibration). Furthermore, Biocrates’ MetIDQ software optionally allows for additional computational batch normalization with respect to mean or median concentrations of repeated “quality control samples” (QC-based normalization). To avoid confusion with nomenclature, we use the term “reference QC” to refer to this standardized quality control (QC) sample provided by Biocrates as part of their kits, and “pooled QC” to refer to our own pooled sample QC.

We utilize multiple measures to assess data quality and to identify discrepancies (Figure 1 – Quality Control). We start with histograms of the number of samples per type, compounds per class as well as measurement statuses (Figure 1 – Status Profiling) as an initial overview of the data set. The number of missing values per sample is visualized via histograms for different sample types to confirm consistent distribution and reasonable superiority of QC samples in terms of detection rate. Frequencies of missing values per compound are visualized with respect to class, sample type and even study groups (if known) to assess differences in occurrences. Sample measurements (such as concentration, intensity, area, etc.) are summarized to establish general sample behavior in (multi-) batch contexts and in particular to identify systematic errors: Visualizing total sample concentration (or missing values) with respect to acquisition order (Figure 1 – Sequence Progression) enables the identification of temporal errors and trends such as batch drifts and carryovers, confirmation of potential corrections such as IS- and QC-based normalization and calibration as well as systematic comparison of batches.

**Figure 1:**
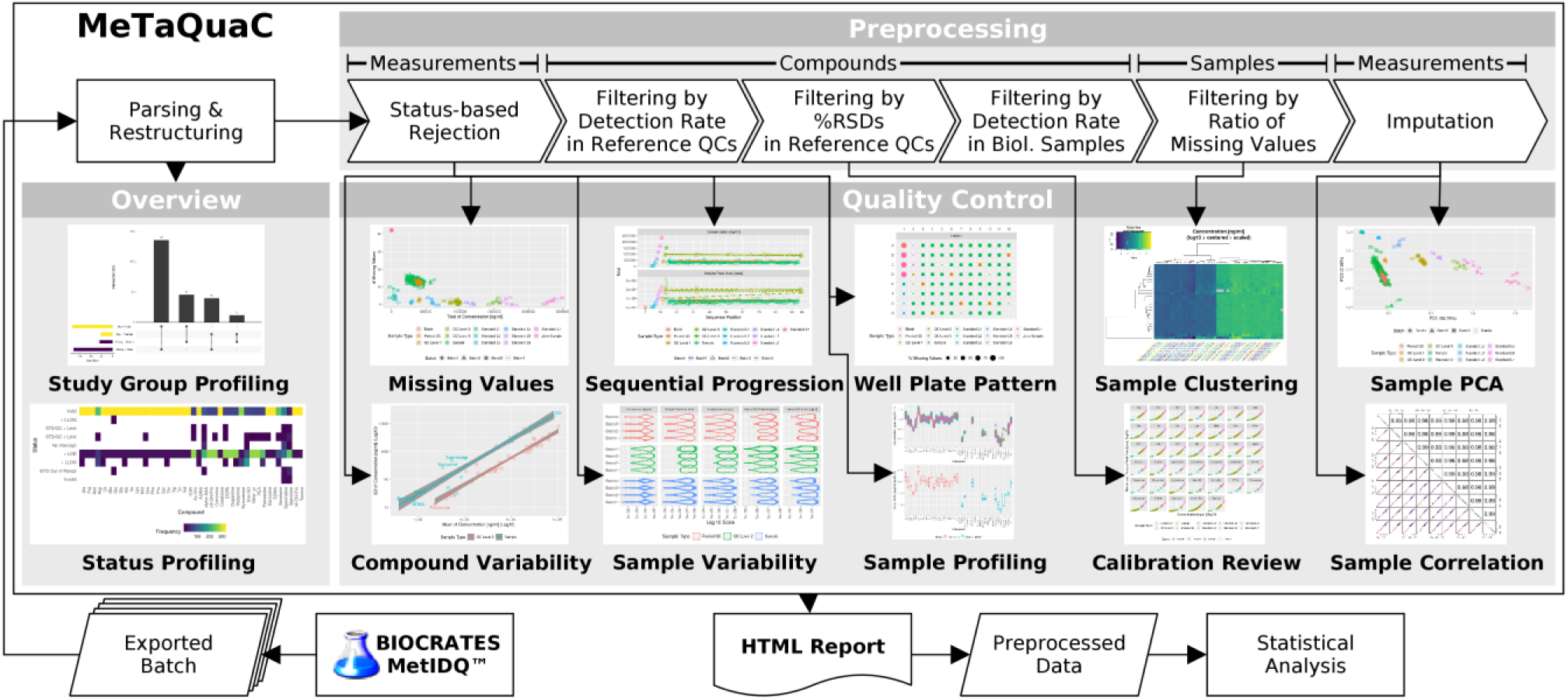
MeTaQuaC quality control workflow. The workflow illustrates the main stages of MeTaQuaC including data import, configurable preprocessing, overview and quality control. It only features a representative extract of overview and quality visualizations, with many more being available in the actual report.

Statistical analysis of linear model coefficients supports the evaluation of drifts within batches and differences between batches. Compound variability in the form of (relative) standard deviation plotted with respect to average concentrations (Figure 1 – Compound Variability) enables confirmation of the essential separation between technical and biological variance. Additional statistical analysis of linear model coefficients is used to confirm the separation. Finally, unsupervised multivariate analysis such as hierarchical clustering, correlation and principal component analysis (Figure 1 – Sample Clustering, Sample Correlation and Sample PCA) represent additional informative methods to further validate conformity within and differentiation between different sample types as well as to provide preliminary insight into study variables and groups.

In addition to and in preparation for QC analysis, our workflow applies several optional but recommended preprocessing steps (in the described order and as shown in Figure 1). First, unreliable measurements are discarded based on Biocrates status flags (as default, only “Valid” concentrations are retained) and considered as missing values (“Status-based Rejection”). Then, unreliable compounds may be removed based on the ratio of missing values (“Filtering by Detection Rate in Reference QCs”) and relative standard deviation in QC samples (kits’ QC Level 2, “Filtering by %RSD in Reference QCs”), while underrepresented compounds may be filtered by ratio of missing values in biological samples (“Filtering by Detection Rate in Biol. Samples”), with defaults of 30%, 15% and 30%, resp., according to Broadhurst et al. ^5^. Furthermore, missing value imputation based on median may be applied but is only recommended after rigorous filtering. In general, subsequent analyses do not always rely on all preprocessing steps (Figure 1 but as many as technically necessary (e.g. a PCA requires a complete dataset and thus requires full preprocessing).

## Results and Discussion

We designed the MeTaQuaC R package to implement the proposed QC and processing measures, among others, for targeted data acquired using Biocrates kits and present them in an extensive HTML report based on R Markdown. Figure 1 depicts the QC workflow for processing targeted metabolomics data as implemented with MeTaQuaC. It includes a parser for Biocrates’ MetIDQ data, data restructuring including batch merging, overview visualizations, data preprocessing including filtering and various quality control visualization and evaluations. Restructured, preprocessed and other intermediary data may be accessed from the report (as csv) and can be exported for further statistical analysis. MeTaQuaC currently supports Biocrates’ AbsoluteIDQ® p400 HR Kit and MxP® Quant 500 Kit.

We demonstrate the application of the proposed measures and of the MeTaQuaC software on two genuine but anonymized data sets. Data set one consists of 240 biological samples and was measured with Biocrates AbsoluteIDQ p400 HR Kit on an Agilent 1290 UHPLC Infinity column and Thermo Fischer Q Exactive Plus mass spectrometer in four batches. The results have been processed with the Biocrates’ MetIDQ software version 7.11.5-DB108-Nitrogen-2834, and the corresponding data files can be found in the supplement (files: data_p400.zip). The complete MeTaQuaC reports are also provided as supplement (file: qc_p400.zip). Data set two consists of eight biological samples and was measured with Biocrates MxP Quant 500 Kit on an Agilent 1290 UHPLC Infinity column and Sciex 5500 mass spectrometer in one batch. The results have been processed with the Biocrates MetIDQ software version 7.11.5-DB108-Nitrogen-2834, and the corresponding data files can be found in the supplement (file: data_q500.zip). The complete MeTaQuaC reports are also provided as supplement (file: qc_q500.zip). Each report contains a multitude of different plots, interactive tables and statistical measures. The computation and generation of the reports takes just a few minutes on a modern desktop computer.

We highlight the benefit of proposed quality control measures with two exemplary visualizations from the analysis of data set one: Figure 2 compares the concentration and total area of LC measurements of each sample in acquisition sequence order. This visualization allows the inspection of several key characteristics of the data: First, it demonstrates the overall total effect of normalizations and calibration. A successful normalization and calibration will result in total concentrations of all QC samples to be on the same level. This leads to horizontal regression lines in the plot (Biocrates provided QC Level 2 samples = green; User-provided pooled samples = orange, if available). Intra-batch differences show as deviations from the horizontal line and are easily spotted in the plot as well as quantitatively described by regression model parameter given in the report. Second, the same feature directly indicates whether different batches are comparable (as expected) or subject to major discrepancies (inter-batch differences visualized by offsets between the regression lines and quantified by the regression coefficients). Another way of double checking normalization performance is provided by the visualization of the standard samples (left hand side of plots in Figure 2). A successful normalization maps standards with equal concentrations closer to each other (with typical higher variance in higher concentrated samples) and clearly separates them from other concentrations. Finally, strong single outliers may be easily spotted (here none). For the data set at hand we conclude satisfactory calibration and normalization performance.

**Figure 2:**
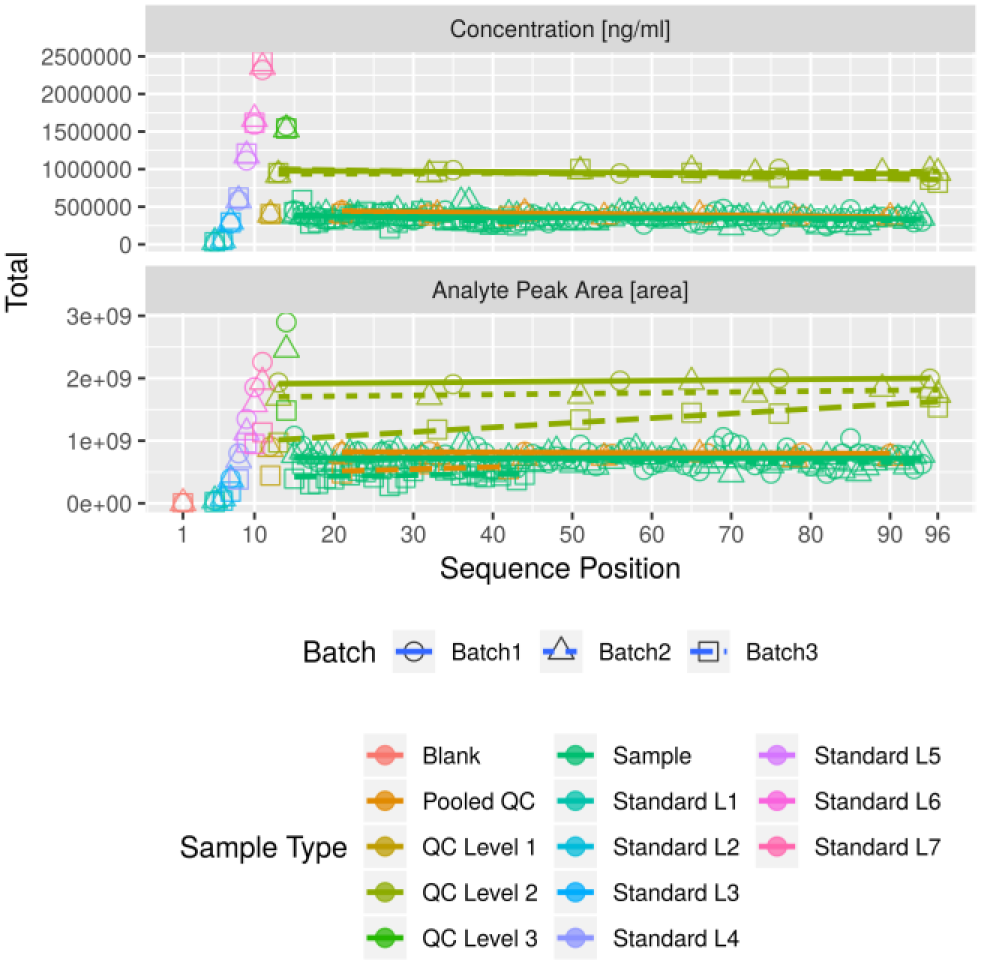
Selected visualization from MeTaQuaC report for data set one comparing intra- and inter-batch consistency of LC data using total analyte peak area (lower plot) and total concentration (upper plot). Data is plotted in sequence of acquisition to visualize and identify potential intra-batch drift and validate normalization and calibration.

Figure 3 visualizes the occurrence of missing values of FIA measurements in each sample with respect to well plate position. It allows for an easy detection of patterns from spatially adjacent samples, which are hard to recognize in the sequential visualization. In this example, a slight increase of missing values can be observed in the connected samples at row A, column 3 to 7 in batch one, indicating an edge effect (e.g. due to lateral drying). Furthermore, the visualization confirms the expected low number of missing values in kit-based QC samples as well as the expected high number in blank and zero samples.

**Figure 3:**
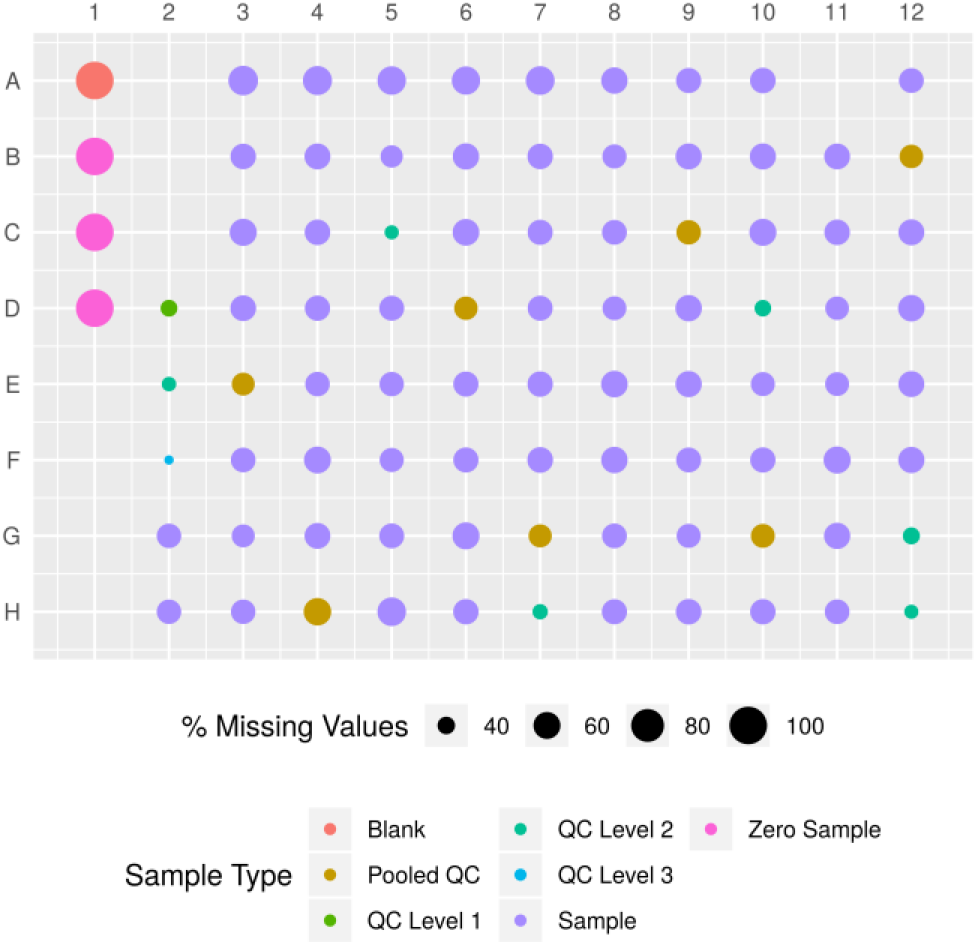
Selected visualization from MeTaQuaC report for data set one comparing spatial distribution of missing values across the well plate for FIA data (hence no calibration standard samples). Samples are plotted per well plate position to visualize positional pattern.

Applying a host of different visualizations and statistics enables changes in perspective and thus often helps to confirm observed conditions, to validate assumptions or to detect or highlight distinct issues. To this end, MeTaQuaC implements various QC measures beyond the once presented. The highlighted examples only cover a small selection of the provide QC measures, but clearly demonstrate the possibility to quickly and efficiently detect a wide range of complex systematic errors and outliers. Furthermore they may aid validating critical processes such as calibration or normalizations. While we compute and visualize many different QC measures and reject measurements, compounds and samples based on user defined thresholds for further downstream QC and potentially statistical analysis, we intentionally do not suggest any thresholds for rejecting or accepting individual batches or whole data sets. To our experience these thresholds depend critically on the individual study, e.g. exploratory versus confirmatory. However, we encourage labs and individual users to select and define their own set of most informative visualizations, measures and acceptance ranges for the different use cases in targeted metabolomics.

## Conclusions

The increasing amounts of studies and data due to advancements in technology demands a shift towards automated and thus robust, reproducible and reportable quality control. The metabolomics community has already recognized the need for more quality assurance and control in metabolomics studies. Community-wide collaborations such as the Metabolomics Quality Assurance and Quality Control Consortium (mQACC) ^21^ further highlight the need and interest in quality control by creating a forum to discuss and implement community-wide procedures and standards for quality assurance and quality control. mQACC is providing education and is developing standard reference material with focus on discovery-based (i.e. untargeted) methods. Our software complements these efforts with easy to apply measures for target studies in a readily usable, consistent and reproducible manner. Although currently limited to Biocrates kits, it can be generalized for generic targeted metabolomics data in the future. MeTaQuaC is available as an open source R package under a permissive MIT license. Source code, documentation and issue tracking can be found at https://github.com/bihealth/metaquac.

## ASSOCIATED CONTENT

### Supporting Information

Data measured with AbsoluteIDQ® p400 HR Kit (data_p400.zip)

MeTaQuaC report for AbsoluteIDQ® p400 HR Kit data (qc_p400.zip)

Data measured with MxP® Quant 500 Kit (data_q500.zip)

MeTaQuaC report for MxP® Quant 500 Kit data (qc_q500.zip)

## AUTHOR INFORMATION

### Author Contributions

DB and JK devised the study and manuscript, MK build the workflow, created the software and worked on the manuscript, NK, DL, VS and AE provided samples and generated data. All authors have given approval to the final version of the manuscript.

### Ethics and Animal Welfare

Ethical approval was obtained from the local ethics committee (No.EA1/270/16) and the study was conducted in accordance with the Declaration of Helsinki. Written informed consent was obtained from all participating subjects and the study was registered at ClinicalTrials.gov (NCT03129503). All animal experimentation was performed in accordance with institutional guidelines following approval by the local authorities of the Federal State of Berlin (X9007/17).

## ACKNOWLEDGMENT

We acknowledge fruitful discussion of the QC workflow and report with members of the Metabolomics and Bioinformatics Core Units. Various members of the Bioinformatics Core Unit contributed code snippets for specific plots.

