## Supplementary material for "Concepts and Software Package for Efficient Quality Control in Targeted Metabolomics Studies – MeTaQuaC": QC p400: qc_p400_1_lc.nb.html

Biocrates QC - p400 - LC


Code 

- Show All Code
- Hide All Code
- Download Rmd

### Biocrates QC - p400 - LC

###### *mkuhring*

###### *January 08, 2020*


This report was created with MeTaQuaC v0.1.1.

The data as imported, restructured and preprocessed in this report is available for export in the section Preprocessing (via the full data tables provided for each preprocessing step).

Before reading and interpreting this QC report, please make sure to be familiar with the Biocrates kit used, i.e. familiarize your self with the compounds, sample types, status values, terminology, analytical specification, etc.  
Please refer to Biocrates’ manuals and documents provided with the kit used.

### 1 Data Preparation


#### 1.1 Import

Import of **Biocrates AbsoluteIDQ p400 HR Kit** data of measurement type **LC**.

Data files to import:


- **Batch1**: */data/mkuhring/workspace/metaquac/inst/extdata/biocrates\_p400\_test\_01/Batch1\_LC1.txt* and */data/mkuhring/workspace/metaquac/inst/extdata/biocrates\_p400\_test\_01/Batch1\_LC2.txt*
- **Batch2**: */data/mkuhring/workspace/metaquac/inst/extdata/biocrates\_p400\_test\_01/Batch2\_LC1.txt* and */data/mkuhring/workspace/metaquac/inst/extdata/biocrates\_p400\_test\_01/Batch2\_LC2.txt*
- **Batch3**: */data/mkuhring/workspace/metaquac/inst/extdata/biocrates\_p400\_test\_01/Batch3\_LC1.txt* and */data/mkuhring/workspace/metaquac/inst/extdata/biocrates\_p400\_test\_01/Batch3\_LC2.txt*


Importing batch `Batch1`…

Importing batch `Batch2`…

Importing batch `Batch3`…

#### 1.2 Restructuring

The imported data was restructured as follows:

- Convert to **long table** (or molten dataset). Each row includes only one measurement, i.e. the data of one compound in one sample and all corresponding information.
- Add **Sample Identification “Blank”** for blank samples, as they are empty when exported from MetIDQ.
- Add **Sample Type “Pooled QC”** for pooled QC samples. Pooled QC samples are identified by any occurrence of the term “pool” (case-insensitive) in the column Sample.Identification (parameter `pool_indicator`). In contrast, **“Reference QC”** will refer to Biocrates’ QC Level 2 samples.
- Add **Sequence Position**, i.e. sample acquisition order. This position is calculated from MetIDQ’s Well Position, as this one depicts a position by row-major order which does not reflect the actual acquisition order.
- Add **Well Coordinates**, indicating the two dimensional position on the well plate. This position is calculated from MetIDQ’s Well Position and indicates rows by letters and columns by numbers.
- Add a **unique Sample Name**, which is a combination of Sample Identifier, Sequence Position and potentially Batch (if available). These names assure unambiguous identification and analysis of measurements within or between several batches (e.g. in particular technical samples such as calibration standard or reference QC samples share sample identifications).
- Remove **nonessential columns**, as some of them currently interfere with proper data merging or processing later. This includes (for now, to be optimized/reduced): Plate Bar Code, Sample Bar Code, Submission Name, Material, Plate Production No., Plate Note, Run Number, Injection Number, Measurement Time, Sample Description, Collection Date, Org. Info and OP.
- Unify **missing values**, i.e. “” (empty field), “0” (zero) and “NA” (not available) are converted to “NA”.
- **Factorize Compounds** with level order based on compound class and compound name.

### 2 Overview


#### 2.1 Samples and Compounds


##### 2.1.1 Samples


**Number of samples in total:** 240

**List of all samples:**


**Number of samples per batch:**

##### 2.1.2 Compounds


**Number of compounds in total:** 42

**List of all compounds:**


**Number of compounds per batch:**

#### 2.2 Study Variables Profile


This study variables profile visualizes group sizes within study variables as well as intersected with other study variables, if more than one is indicated. Set Size = Number of samples per group in variable (color-coded). Intersection Size = Number of samples with same combination of groups.The following variable(s) are used: **Group**, **Condition**.

#### 2.3 Status Profiles

Status profiles visualize the occurrence of the different possible measurement statuses. Profiles are generated over all measurements, for different sample types (in percentages, since the number of samples vary per type), for different samples, for different compound classes (in percentages, since the number of compounds vary per class), as well as for different compounds.


##### 2.3.1 Overall

##### 2.3.2 Per sample type


###### 2.3.2.1 Heatmap

###### 2.3.2.2 Single

##### 2.3.3 Per sample


###### 2.3.3.1 Heatmap

###### 2.3.3.2 Single

##### 2.3.4 Per compound class


###### 2.3.4.1 Heatmap

###### 2.3.4.2 Single

##### 2.3.5 Per compound


###### 2.3.5.1 Heatmap

###### 2.3.5.2 Single

### 3 Preprocessing


The following preprocessing procedures are applied on the data in this order:

1. **Status Preprocessing**
   1. Discard unreliable measurements based on Biocratus status
2. **Remove unreliable **compounds**** (QC-based filters)
   1. Based on missing values ratio in Reference QCs (**>=30%**)
   2. Based on %RSD of Reference QCs (**>=15%**)
3. **Remove underrepresented or invariable **compounds**** (biological filters)
   1. Based on only missing values in biological samples (**=100%**)
   2. Based on missing values ratio in biological samples (**>=30%**)
   3. Based on %RSD in biological samples (**<15%**)
4. **Remove underrepresented **biological samples**** (biological filters)
   1. Based on missing values ratio of compounds (**>=20%**)

The preprocessing statuses and filter thresholds can be modified with the following parameters of the `create_report` function (refer to the readme or function description for details):

- `preproc_keep_status`
- `filter_compound_qc_max_mv_ratio`
- `filter_compound_qc_max_rsd`
- `filter_compound_bs_max_mv_ratio`
- `filter_compound_bs_min_rsd`
- `filter_sample_max_mv_ratio`

The result of each preprocessing or filter step can be found in the tab with the corresponding enumerator. Please note, as each tab contains a table with the complete dataset in the corresponding filtered form, loading the tab for the first time might take a couple of seconds.


**Note**: The beginning of each following section will indicate on which dataset calculations and visualizations are based. Thereby, the preprocessed datasets will be referenced as **1**, **2a**, **2b**, **3a**, **3b** and **4**, resp.


#### 3.1 Step **1**

1. **Status Preprocessing**
   1. Discard unreliable measurements based on Biocratus status

The status preprocessing discards measurements based on Biocrates quality statuses.

By default, only “Valid” measurements are considered as reliable and are kept for further analysis. Hence, all other statuses including “< LLOQ” and “> ULOQ” (out of quantification range / extrapolated calibration), “STD/QC < Limit” and “STD/QC > Limit” (insufficient accuracy in STDs and QCs), “ISTD Out of Range” (internal standard too high or low) and other statuses are usually considered as unreliable and measurements are transformed to missing values (set to NA).

Based on the parameter `preproc_keep_status`, measurements with the following statuses will be retained: **Valid**

Missing **Concentration** values before Biocrates status preprocessing: **2565** of **10080** (25%)

Missing **Concentration** values after Biocrates status preprocessing: **3069** of **10080** (30%)

#### 3.2 Step **2a**

2. **Remove unreliable **compounds**** (QC-based filters)
   1. Based on missing values ratio in Reference QCs (**>=30%**)

To alter the threshold use parameter `filter_compound_qc_max_mv_ratio`.

Number of compounds before: **42** Number of compounds left: **39**

#### 3.3 Step **2b**

2. **Remove unreliable **compounds**** (QC-based filters)
   2. Based on %RSD of Reference QCs (**>=15%**)

To alter the threshold use parameter `filter_compound_qc_max_rsd`.

Number of compounds before: **39** Number of compounds left: **39**

#### 3.4 Step **3a**

3. **Remove underrepresented **compounds**** (biological filters)
   1. Based on only missing values in biological samples (**=100%**)

Number of compounds before: **39** Number of compounds left: **30**

#### 3.5 Step **3b**

3. **Remove underrepresented **compounds**** (biological filters)
   2. Based on missing values ratio in biological samples (**>=30%**)

To alter the threshold use parameter `filter_compound_bs_max_mv_ratio`.

Number of compounds before: **30** Number of compounds left: **27**

#### 3.6 Step **3c**

3. **Remove underrepresented **compounds**** (biological filters)
   3. Based on %RSD in biological samples (**<15%**)

To alter the threshold use parameter `filter_compound_bs_min_rsd`.

Number of compounds before: **27** Number of compounds left: **27**

#### 3.7 Step **4**

4. **Remove underrepresented **biological samples**** (biological filters)
   1. Based on missing values ratio of compounds (**>=20%**)

To alter the threshold use parameter `filter_sample_max_mv_ratio`.

Number of all samples before: **240** Number of all samples left: **228**

Number of biological samples before: **164** Number of biological samples left: **164**

#### 3.8 Original

Missing value counts either by samples or by compounds based on unprocessed data (i.e. no values have been turned into missing values based on status and nothing is filtered) as well as the full unprocessed dataset is available in the tables below.

### 4 Quality Control


The following QC analyses are based on the status-preprocessed dataset 1, unless explicitly stated otherwise.


#### 4.1 Missing values

##### 4.1.1 Per sample

Missing values are counted per sample, i.e. how many compounds were not measured in the sample at all or reliably enough (depending on status preprocessing). First, missing value counts are visualized depending on the total concentration measured in a sample (with Well Positions indicated for a few data points in low density areas).


Furthermore, missing value counts are summarized by histograms (primarily) including visual separation and comparison of relevant sample types, batches (if applicable) or study variable groups (if indicated).

###### 4.1.1.1 Per sample type

###### 4.1.1.2 In batches

###### 4.1.1.2.1 All samples

###### 4.1.1.2.2 Reference QCs

###### 4.1.1.2.3 Pooled QCs

###### 4.1.1.2.4 Biological sample

###### 4.1.1.3 Per study variable

###### 4.1.1.3.1 Group

###### 4.1.1.3.2 Condition

###### 4.1.1.3.3 Group = Control -> Condition

###### 4.1.1.3.4 Group = Case -> Condition

##### 4.1.2 Per compound

Missing values are counted per compound, i.e. how many samples don’t feature a reliable measurement of the compound (depending on status preprocessing). Counts are calculated and visualized for different compound classes as well as for different sample subsets. I.e., for the same compound, missing values are separately counted within different sample types or within different study variable groups (if indicated).

###### 4.1.2.1 Per compound class

###### 4.1.2.2 Per sample type

###### 4.1.2.3 Per study variable

Bar plots per study variable visualize differences in the occurrence of missing values in different study groups. **# Missing Values** counts are used as reference to show whether a compound is missing just a bit or a lot. It shouldn’t be used for comparison of groups due to possible differences in group sizes. However, **% Missing Values** show the percentage of missing values per group, which are further added up and normalized to one as **Normalized % Missing Values**. This allows to infer compounds which are considerably more missing in certain groups (if the missing value count is high). Only compounds with a least one and less than 100% missing values (overall) are shown.

###### 4.1.2.3.1 Group

###### 4.1.2.3.2 Condition

###### 4.1.2.3.3 Group = Control -> Condition

###### 4.1.2.3.4 Group = Case -> Condition

#### 4.2 Sample variability

##### 4.2.1 Sequence progression

Progression of sample summary statistics (totals for concentration/area/intensity and missing values) is visualized over the acquisition sequence. This allows an quick overview of variability between replicate samples as well as between batches, while the parallel view on the different data types (concentration, area, istd area, etc.) illustrates the effect of calibration/normalization.

After calibration (concentration), replicate samples (such as reference and pooled QCs) should feature a preferably horizontal regression line (in particular for primarily multi-point calibrated data). Furthermore, regression lines of different batches should align reasonably to justify subsequent joint analysis. In comparison, uncalibrated and unnormalized measurements (area and intensity) may feature incomparable fluctuation and batch drifts.


###### 4.2.1.1 Totals overview

###### 4.2.1.2 Totals details

Linear models are calculated to assess the horizontal behavior of measurements (total concentration per sample) within specific sample types. In particular, in a perfect batch with no deviation the resulting regression lines should be horizontal. Statistical analysis is applied to highlight potential deviation of the regression line fitted to the experimental data from the horizontal.

For **single batches**, the simple linear model `Total Concentration ~ Sequence Position` is checked for unwanted batch drifts inferred from significant slopes. Significant slopes may not only indicate batch drifts but general technical variability not sufficiently resolved by internal standard normalization and external standard calibration.

For **batch comparison**, the model `Total Concentration ~ Batch + Sequence Position / Batch` is used to infer unwanted significant differences in intercepts and slopes between batches (with one batch randomly selected as reference). Furthermore, a Kruskal-Wallis rank sum test is used to infer unwanted general differences in batch distributions.


**Note**: Use the automated rating for guidance only and in addition judge manually (e.g. by visual inspection of the totals overview) whether total concentration fits behave well enough in comparison to area or intensity (i.e. unnormalized and uncalibrated data). The rating remains rather subjective for now, as it will require a great number of experiments and extensive validation to assess which model coefficients and p-values are acceptable to support reliable analysis.


###### 4.2.1.2.1 Reference QCs


###### 4.2.1.2.1.1 Single batches

No batch with undesirable Concentration [ng/ml] slope for samples of type QC Level 2.

###### 4.2.1.2.1.2 Batch comparison

**Warning:** Detected significant differences in batch slopes of total Concentration [ng/ml] for samples of type QC Level 2 (delta p-value < 0.01)!

Kruskal-Wallis p-value: 0.536327549854899

Model fit: 0.55567532401262

###### 4.2.1.2.1.3 Auxiliary Figures

###### 4.2.1.2.2 Pooled QCs

###### 4.2.1.2.2.1 Single batches

**Warning:** Batch **Batch1** features a significant Concentration [ng/ml] slope for samples of type Pooled QC (p-value < 0.01)!

###### 4.2.1.2.2.2 Batch comparison

There are no significant differences in total Concentration [ng/ml] between batches for samples of type Pooled QC.

Kruskal-Wallis p-value: 0.472918189432697

Model fit: 0.760489333622129

###### 4.2.1.2.2.3 Auxiliary Figures

###### 4.2.1.2.3 Biological samples


###### 4.2.1.2.3.1 Single batches

No batch with undesirable Concentration [ng/ml] slope for samples of type Sample.

###### 4.2.1.2.3.2 Batch comparison

There are no significant differences in total Concentration [ng/ml] between batches for samples of type Sample.

Kruskal-Wallis p-value: 0.356960783225124

Model fit: 0.094395813062693

###### 4.2.1.2.3.3 Auxiliary Figures

###### 4.2.1.3 Missing values overview

##### 4.2.2 Well Plate Overview

Sample summary statistics (missing values and total concentration/area/intensity) are compared with respect to well plate position. This allows to identify unexpected position-based issues (e.g. edge-effects), as might be demonstrated by a cluster of aberrating dots, or to spot outlier samples. In general, samples of the same type should feature similar dot sizes (with minor variations in biological samples). Samples areas and intensities will vary more than concentrations (as latter are calibrated).


###### 4.2.2.1 % Missing Values of Concentration

###### 4.2.2.2 Total Concentration

###### 4.2.2.3 Total Area

###### 4.2.2.4 Total Intensity

###### 4.2.2.5 Total Area ISTD

###### 4.2.2.6 Total Intensity ISTD

##### 4.2.3 Samples overlapping by type

Violin and box plots summarize compound measurements per sample and are grouped and overlapped per sample type, if applicable. This allows an quick overview of variability between replicate samples such as reference and pooled QCs as well as biological samples, while the parallel view on the different data types illustrates the effect of calibration/normalization (i.e. replicate sample concentration profiles should line up rather well).


###### 4.2.3.1 Violins

###### 4.2.3.2 Boxplots

##### 4.2.4 Samples ordered by aquisition

Violin and box plots summarize compound measurements per replicate sample, grouped per sample type and ordered according to acquisition sequence. This allows a quick overview of variability progression of replicate samples such as QCs or Pool samples over the batch (e.g. to indicate batch drift), while the parallel view on the different data types illustrates the effect of calibration/normalization (i.e. replicate sample concentration profiles should line up rather well).


###### 4.2.4.1 Violins

###### 4.2.4.2 Boxplot

#### 4.3 Compound variability

##### 4.3.1 Overview

The plot below shows the standard deviation of each individual metabolite against their average concentration, separately for QC and biological samples (i.e. technical vs biological variability).

A linear regression analysis is used to attempt to separate the technical and biological variability. The outputs of the models are used to score and assess this separation, and can be used as indicator for data quality.


There is a clear separation between technical and biological variability.


Please see the details and the plot below for more information.

##### 4.3.2 Details

An integrative linear model is used to assess differences in technical and biological variability: `log(SD) ~ Group + log(Mean)/Group` with Group being a factor indicating QC or biological samples.

Model results are combined to identify four important main cases of differences in technical and biological variability:

1. `k >= 7`: A clear biological variability on top of the technical one.
2. `k = 2:6`: Inconclusive differences in biological and technical variability.
3. `k = 0:1`: A strong technical variability which hides the biological one.
4. `k < 0`: Something clearly wrong (when variability in QC decreases with increasing amount of metabolite concentration).

with final score k being computed to allow the classification of the four main cases as follows:

```
k = 0
k = k + (r_squared > 0.8)
k = k + 2*(delta_intercept > 0 & delta_intercept_pvalue < 0.05)
k = k + 4*(delta_slope > 0)
k = k - 8*(qc_slope < 0)
```

The results are as follows:

- Linear regression model results:
  - R squared = 0.9733469
  - QC slope = 0.9519574
  - Delta slope (Sample - QC) = 0.0079493
  - Delta slope p-value = 0.8394102
  - Delta intercept (Sample - QC) = 1.46615
  - Delta intercept p-value = 8.491990210^{-5}
- Variability Score:
  - k = 7

##### 4.3.3 Per study variable

For visual examination, SD as well as %RSDs are plotted against the mean per compound not only separated by sample type but further separated by groups in study variables of the biological samples.

###### 4.3.3.1 Group

###### 4.3.3.2 Condition

###### 4.3.3.3 Group = Control -> Condition

###### 4.3.3.4 Group = Case -> Condition

#### 4.4 Compound %RSDs

Relative standard deviations per compound or compound class are calculated and visualized for different sample groupings, either by sample type or for biological replicates (i.e. samples from the same group within variables indicated with parameter `replicate_variables`) as well as for all batches or each batch separately.

This analysis is based on the status-preprocessed dataset 1.


##### 4.4.1 Compounds

###### 4.4.1.1 All batches


###### 4.4.1.1.1 Sample types

###### 4.4.1.1.2 Study groups (separate)

###### 4.4.1.1.3 Study groups (interacting)

###### 4.4.1.2 Per batch


###### 4.4.1.2.1 Sample types

###### 4.4.1.2.2 Study groups (separate)

###### 4.4.1.2.3 Study groups (interacting)

##### 4.4.2 Class medians

###### 4.4.2.1 All batches

###### 4.4.2.1.1 Sample types

###### 4.4.2.1.2 Study groups (separate)

###### 4.4.2.1.3 Study groups (interacting)

###### 4.4.2.2 Per batch

###### 4.4.2.2.1 Sample types

###### 4.4.2.2.2 Study groups (separate)

###### 4.4.2.2.3 Study groups (interacting)

##### 4.4.3 Sample type medians

###### 4.4.3.1 All batches

###### 4.4.3.2 Per batch

#### 4.5 Calibration

Calibration scatter plots are reconstructed to enable an impression of calibration performance and range placement (in particular for 7-point calibrated compounds). To properly reflect the internal standard calibration method, areas and intensities are normalized by the ISTD areas and intensities, resp.

The following visualization is based on the status-preprocessed dataset 2b.


##### 4.5.1 Concentration vs peak area

##### 4.5.2 Concentration vs intensity

### 5 Multivariate Visualization


The following multivariate analyses are based on the preprocessed dataset 4, unless explicitly stated otherwise.


#### 5.1 Clustered heatmaps

Heatmaps of centered and scaled log10 target values (i.e. concentration, intensity or area) by compounds and samples. Sample labels may be colored by additional variables (e.g. Batch or Sample.Type). Compounds as well as samples are hierarchically clustered with Pearson correlation as distance metric or euclidean distance metric (aka L2 norm).


##### 5.1.1 Pearson

###### 5.1.1.1 All sample types


###### 5.1.1.1.1 Batch-colored labels

###### 5.1.1.1.2 Type-colored labels

###### 5.1.1.2 QC samples


###### 5.1.1.2.1 Batch-colored labels

###### 5.1.1.2.2 Type-colored labels

###### 5.1.1.3 Biological samples


###### 5.1.1.3.1 Batch-colored labels

###### 5.1.1.3.2 Group-colored labels

###### 5.1.1.3.3 Condition-colored labels

###### 5.1.1.3.4 Group = Control -> Condition-colored labels

###### 5.1.1.3.5 Group = Case -> Condition-colored labels

##### 5.1.2 Euclidean

###### 5.1.2.1 All sample types


###### 5.1.2.1.1 Batch-colored labels

###### 5.1.2.1.2 Type-colored labels

###### 5.1.2.2 QC samples


###### 5.1.2.2.1 Batch-colored labels

###### 5.1.2.2.2 Type-colored labels

###### 5.1.2.3 Study samples


###### 5.1.2.3.1 Batch-colored labels

###### 5.1.2.3.2 Group-colored labels

###### 5.1.2.3.3 Condition-colored labels

###### 5.1.2.3.4 Group = Control -> Condition-colored labels

###### 5.1.2.3.5 Group = Case -> Condition-colored labels

#### 5.2 Missing value imputation

Median imputation is applied for the remaining visualizations on biological, reference and pooled QC samples (separately), as the methods are not suited to handle missing values. Samples which still contain missing values after imputation are completely removed.

#### 5.3 Sample correlation

Scatter plot matrices illustrate the correlation between samples based on compound concentration.

In the case of high sample numbers (> 9), several matrices will be generated based on sample groups of size up to 9 with one overlapping sample between each group (i.e. the last sample in a matrix is the first sample in the next matrix).

This analysis is based on the imputed dataset 4.

##### 5.3.1 Reference QCs


#### 5.3.1.1 1

#### 5.3.1.2 2

#### 5.3.1.3 3

##### 5.3.2 Pooled QCs

#### 5.3.2.1 1

#### 5.3.2.2 2

#### 5.3.2.3 3

##### 5.3.3 All samples


#### 5.3.3.1 1

#### 5.3.3.2 2

#### 5.3.3.3 3

#### 5.3.3.4 4

#### 5.3.3.5 5

#### 5.3.3.6 6

#### 5.3.3.7 7

#### 5.3.3.8 8

#### 5.3.3.9 9

#### 5.3.3.10 10

#### 5.3.3.11 11

#### 5.3.3.12 12

#### 5.3.3.13 13

#### 5.3.3.14 14

#### 5.3.3.15 15

#### 5.3.3.16 16

#### 5.3.3.17 17

#### 5.3.3.18 18

#### 5.3.3.19 19

#### 5.3.3.20 20

#### 5.3.3.21 21

#### 5.3.3.22 22

#### 5.3.3.23 23

#### 5.3.3.24 24

#### 5.3.3.25 25

#### 5.3.3.26 26

#### 5.3.3.27 27

#### 5.3.3.28 28

#### 5.3.3.29 29

#### 5.4 PCA

Principal component analysis (PCA) is performed to emphasize compound concentrations-based sample type, batch and potential study group relationships.

This analysis is based on the imputed dataset 4.

##### 5.4.1 All samples

##### 5.4.2 Biological vs QC samples

##### 5.4.3 Biological samples (colored groups)

###### 5.4.3.1 Group

###### 5.4.3.2 Condition

###### 5.4.3.3 Group = Control -> Condition

###### 5.4.3.4 Group = Case -> Condition

### 6 Miscellaneous


#### 6.1 Additional figures

##### 6.1.1 Replicate dispersion

The following figures illustrate reproducibility by comparing compound measures (e.g. concentration) within and between groups such as different sample types and conditions in study variables (as indicated with parameter `study_variables`).

This analysis is based on status-preprocessed dataset 3a.


###### 6.1.1.1 Per sample type

###### 6.1.1.2 Per study variable


###### 6.1.1.2.1 Group

###### 6.1.1.2.2 Condition

###### 6.1.1.2.3 Group -> Condition

##### 6.1.2 Concentration Profiles

Simple metabolite concentration profiles overlapped per sample (point and line plot) or summarized (as box plots) are created to enable a brief impression of overall sample behavior consistency (e.g. to justify later normalization).

This analysis is based on the status-preprocessed dataset 1.


###### 6.1.2.1 Reference QCs

###### 6.1.2.2 Pooled QCs

###### 6.1.2.3 Biological samples

#### 6.2 Evironment info


MeTaQuaC package version: 0.1.1

**R version 3.4.4 (2018-03-15)**

\*\*Platform:\*\* x86\_64-pc-linux-gnu (64-bit)

**locale:** *LC\_CTYPE=en\_US.UTF-8*, *LC\_NUMERIC=C*, *LC\_TIME=en\_US.UTF-8*, *LC\_COLLATE=en\_US.UTF-8*, *LC\_MONETARY=en\_US.UTF-8*, *LC\_MESSAGES=en\_US.UTF-8*, *LC\_PAPER=en\_US.UTF-8*, *LC\_NAME=C*, *LC\_ADDRESS=C*, *LC\_TELEPHONE=C*, *LC\_MEASUREMENT=en\_US.UTF-8* and *LC\_IDENTIFICATION=C*

**attached base packages:** *stats*, *graphics*, *grDevices*, *utils*, *datasets*, *methods* and *base*

**other attached packages:** *metaquac(v.0.1.1)* and *testthat(v.2.2.1)*

**loaded via a namespace (and not attached):** *bitops(v.1.0-6)*, *fs(v.1.3.1)*, *xts(v.0.11-1)*, *usethis(v.1.5.1)*, *devtools(v.2.2.1)*, *rprojroot(v.1.3-2)*, *UpSetR(v.1.3.3)*, *tools(v.3.4.4)*, *backports(v.1.1.4)*, *R6(v.2.4.0)*, *DT(v.0.4)*, *rpart(v.4.1-15)*, *KernSmooth(v.2.23-15)*, *DBI(v.1.0.0)*, *lazyeval(v.0.2.2)*, *colorspace(v.1.4-1)*, *withr(v.2.1.2)*, *tidyselect(v.0.2.5)*, *gridExtra(v.2.3)*, *prettyunits(v.1.0.2)*, *processx(v.3.4.1)*, *curl(v.3.3)*, *compiler(v.3.4.4)*, *cli(v.1.1.0)*, *desc(v.1.2.0)*, *labeling(v.0.3)*, *caTools(v.1.17.1.1)*, *scales(v.1.0.0)*, *readr(v.1.1.1)*, *callr(v.3.3.2)*, *stringr(v.1.3.1)*, *digest(v.0.6.18)*, *rmarkdown(v.1.10)*, *R.utils(v.2.9.0)*, *DMwR2(v.0.0.2)*, *base64enc(v.0.1-3)*, *pkgconfig(v.2.0.2)*, *htmltools(v.0.3.6)*, *sessioninfo(v.1.1.1)*, *htmlwidgets(v.1.3)*, *rlang(v.0.4.1)*, *TTR(v.0.23-4)*, *rstudioapi(v.0.8)*, *quantmod(v.0.4-13)*, *shiny(v.1.1.0)*, *zoo(v.1.8-4)*, *jsonlite(v.1.6)*, *crosstalk(v.1.0.0)*, *gtools(v.3.8.1)*, *dplyr(v.0.8.3)*, *R.oo(v.1.22.0)*, *magrittr(v.1.5)*, *Rcpp(v.1.0.3)*, *munsell(v.0.5.0)*, *viridis(v.0.5.1)*, *R.methodsS3(v.1.7.1)*, *stringi(v.1.1.7)*, *yaml(v.2.2.0)*, *ggpmisc(v.0.3.0)*, *MASS(v.7.3-50)*, *pkgbuild(v.1.0.5)*, *gplots(v.3.0.1)*, *plyr(v.1.8.4)*, *grid(v.3.4.4)*, *gdata(v.2.18.0)*, *promises(v.1.0.1)*, *ggrepel(v.0.8.0)*, *crayon(v.1.3.4)*, *lattice(v.0.20-38)*, *pander(v.0.6.3)*, *hms(v.0.4.2)*, *knitr(v.1.25)*, *ps(v.1.3.0)*, *pillar(v.1.4.3)*, *reshape2(v.1.4.3)*, *pkgload(v.1.0.2)*, *glue(v.1.3.1)*, *evaluate(v.0.13)*, *remotes(v.2.1.0)*, *httpuv(v.1.4.5)*, *gtable(v.0.3.0)*, *purrr(v.0.2.5)*, *tidyr(v.0.8.0)*, *assertthat(v.0.2.1)*, *ggplot2(v.3.1.0)*, *xfun(v.0.4)*, *mime(v.0.6)*, *xtable(v.1.8-3)*, *later(v.0.7.5)*, *class(v.7.3-14)*, *viridisLite(v.0.3.0)*, *tibble(v.2.1.3)*, *memoise(v.1.1.0)*, *ellipsis(v.0.3.0)* and *ggfortify(v.0.4.5)*

#### 6.3 Parameters


- **additional\_normalization**: None
- **pqn\_reference\_type**: Sample
- **sample\_filter**:
- **metadata\_extraction**:
- **zero2na**: *TRUE*
- **ignore\_errors**: *TRUE*
- **data\_files**:

  - **Batch1**: */data/mkuhring/workspace/metaquac/inst/extdata/biocrates\_p400\_test\_01/Batch1\_LC1.txt* and */data/mkuhring/workspace/metaquac/inst/extdata/biocrates\_p400\_test\_01/Batch1\_LC2.txt*
  - **Batch2**: */data/mkuhring/workspace/metaquac/inst/extdata/biocrates\_p400\_test\_01/Batch2\_LC1.txt* and */data/mkuhring/workspace/metaquac/inst/extdata/biocrates\_p400\_test\_01/Batch2\_LC2.txt*
  - **Batch3**: */data/mkuhring/workspace/metaquac/inst/extdata/biocrates\_p400\_test\_01/Batch3\_LC1.txt* and */data/mkuhring/workspace/metaquac/inst/extdata/biocrates\_p400\_test\_01/Batch3\_LC2.txt*
- **kit**: Biocrates AbsoluteIDQ p400 HR Kit
- **measurement\_type**: LC
- **title**: Biocrates QC - p400 - LC
- **author**: mkuhring
- **pool\_indicator**: Sample.Identification
- **profiling\_variables**: *Group* and *Condition*
- **study\_variables**:

  - Group
  - Condition
  - **Group**:

    ```
      * Condition
    ```
- **replicate\_variables**: *Group* and *Condition*
- **preproc\_keep\_status**: Valid
- **filter\_compound\_qc\_max\_mv\_ratio**: *0.3*
- **filter\_compound\_qc\_max\_rsd**: *15*
- **filter\_compound\_bs\_max\_mv\_ratio**: *0.3*
- **filter\_compound\_bs\_min\_rsd**: *15*
- **filter\_sample\_max\_mv\_ratio**: *0.2*
- **data\_tables**: all
