## Supplementary material for "Concepts and Software Package for Efficient Quality Control in Targeted Metabolomics Studies – MeTaQuaC": QC Q500: qc_q500_1_lc.nb.html

Biocrates QC - Q500 - LC


Code 

- Show All Code
- Hide All Code
- Download Rmd

### Biocrates QC - Q500 - LC

###### *mkuhring*

###### *January 08, 2020*


This report was created with MeTaQuaC v0.1.1.

The data as imported, restructured and preprocessed in this report is available for export in the section Preprocessing (via the full data tables provided for each preprocessing step).

Before reading and interpreting this QC report, please make sure to be familiar with the Biocrates kit used, i.e. familiarize your self with the compounds, sample types, status values, terminology, analytical specification, etc.  
Please refer to Biocrates’ manuals and documents provided with the kit used.

### 1 Data Preparation


#### 1.1 Import

Import of **Biocrates MxP Quant 500 Kit** data of measurement type **LC**.

Data files to import:


- **Batch1**: /data/mkuhring/workspace/metaquac/inst/extdata/biocrates\_q500\_test\_01/Batch1\_LC.txt


Importing batch `Batch1`…

#### 1.2 Restructuring

The imported data was restructured as follows:

- Convert to **long table** (or molten dataset). Each row includes only one measurement, i.e. the data of one compound in one sample and all corresponding information.
- Add **Sample Identification “Blank”** for blank samples, as they are empty when exported from MetIDQ.
- Add **Sample Type “Pooled QC”** for pooled QC samples. Pooled QC samples are identified by any occurrence of the term “pool” (case-insensitive) in the column Sex (parameter `pool_indicator`). In contrast, **“Reference QC”** will refer to Biocrates’ QC Level 2 samples.
- Add **Sequence Position**, i.e. sample acquisition order. This position is calculated from MetIDQ’s Well Position, as this one depicts a position by row-major order which does not reflect the actual acquisition order.
- Add **Well Coordinates**, indicating the two dimensional position on the well plate. This position is calculated from MetIDQ’s Well Position and indicates rows by letters and columns by numbers.
- Add a **unique Sample Name**, which is a combination of Sample Identifier, Sequence Position and potentially Batch (if available). These names assure unambiguous identification and analysis of measurements within or between several batches (e.g. in particular technical samples such as calibration standard or reference QC samples share sample identifications).
- Remove **nonessential columns**, as some of them currently interfere with proper data merging or processing later. This includes (for now, to be optimized/reduced): Plate Bar Code, Sample Bar Code, Submission Name, Material, Plate Production No., Plate Note, Run Number, Injection Number, Measurement Time, Sample Description, Collection Date, Org. Info and OP.
- Unify **missing values**, i.e. “” (empty field), “0” (zero) and “NA” (not available) are converted to “NA”.
- **Factorize Compounds** with level order based on compound class and compound name.

### 2 Overview


#### 2.1 Samples and Compounds


##### 2.1.1 Samples


**Number of samples in total:** 23

**List of all samples:**


**Number of samples per batch:**

##### 2.1.2 Compounds


**Number of compounds in total:** 106

**List of all compounds:**


**Number of compounds per batch:**

#### 2.2 Study Variables Profile


This study variables profile visualizes group sizes within study variables as well as intersected with other study variables, if more than one is indicated. Set Size = Number of samples per group in variable (color-coded). Intersection Size = Number of samples with same combination of groups.The following variable(s) are used: **Sex**.

geom\_path: Each group consists of only one observation. Do you need to adjust the group aesthetic?

#### 2.3 Status Profiles

Status profiles visualize the occurrence of the different possible measurement statuses. Profiles are generated over all measurements, for different sample types (in percentages, since the number of samples vary per type), for different samples, for different compound classes (in percentages, since the number of compounds vary per class), as well as for different compounds.


**Note**: Many LC compounds in the Biocrates MxP Quant 500 Kit are 1-point calibrated and don’t feature an external standard. Thus, a high number of “< LOD” is to be expected within standard calibration samples.


##### 2.3.1 Overall

Based on the parameter `preproc_keep_status`, measurements with the following statuses will be retained: **Valid**

Missing **Concentration** values before Biocrates status preprocessing: **295** of **2438** (12%)

Missing **Concentration** values after Biocrates status preprocessing: **1296** of **2438** (53%)

#### 3.2 Step **2a**

2. **Remove unreliable **compounds**** (QC-based filters)
   1. Based on missing values ratio in Reference QCs (**>=30%**)

To alter the threshold use parameter `filter_compound_qc_max_mv_ratio`.

Number of compounds before: **106** Number of compounds left: **89**

#### 3.3 Step **2b**

2. **Remove unreliable **compounds**** (QC-based filters)
   2. Based on %RSD of Reference QCs (**>=15%**)

To alter the threshold use parameter `filter_compound_qc_max_rsd`.

Number of compounds before: **89** Number of compounds left: **78**

#### 3.4 Step **3a**

3. **Remove underrepresented **compounds**** (biological filters)
   1. Based on only missing values in biological samples (**=100%**)

Number of compounds before: **78** Number of compounds left: **53**

#### 3.5 Step **3b**

3. **Remove underrepresented **compounds**** (biological filters)
   2. Based on missing values ratio in biological samples (**>=30%**)

To alter the threshold use parameter `filter_compound_bs_max_mv_ratio`.

Number of compounds before: **53** Number of compounds left: **46**

#### 3.6 Step **3c**

3. **Remove underrepresented **compounds**** (biological filters)
   3. Based on %RSD in biological samples (**<15%**)

To alter the threshold use parameter `filter_compound_bs_min_rsd`.

Number of compounds before: **46** Number of compounds left: **33**

#### 3.7 Step **4**

4. **Remove underrepresented **biological samples**** (biological filters)
   1. Based on missing values ratio of compounds (**>=20%**)

To alter the threshold use parameter `filter_sample_max_mv_ratio`.

Number of all samples before: **23** Number of all samples left: **13**

Number of biological samples before: **8** Number of biological samples left: **8**

#### 3.8 Original


**Note**: Many LC compounds in the Biocrates MxP Quant 500 Kit are 1-point calibrated and don’t feature an external standard. Thus, a high number of missing values is to be expected within standard calibration samples after preprocessing according to status (mainly due to “< LOD”). For the same reason, standard calibration samples are not expected to encompass all the other samples with respect to totals of concentration, area or intensity. For instance, the Biocrates QC Level 3 sample should feature higher totals than the highest standard calibration samples.


###### 4.2.1.1 Totals overview

###### 4.2.1.2 Totals details

Linear models are calculated to assess the horizontal behavior of measurements (total concentration per sample) within specific sample types. In particular, in a perfect batch with no deviation the resulting regression lines should be horizontal. Statistical analysis is applied to highlight potential deviation of the regression line fitted to the experimental data from the horizontal.

For **single batches**, the simple linear model `Total Concentration ~ Sequence Position` is checked for unwanted batch drifts inferred from significant slopes. Significant slopes may not only indicate batch drifts but general technical variability not sufficiently resolved by internal standard normalization and external standard calibration.

For **batch comparison**, the model `Total Concentration ~ Batch + Sequence Position / Batch` is used to infer unwanted significant differences in intercepts and slopes between batches (with one batch randomly selected as reference). Furthermore, a Kruskal-Wallis rank sum test is used to infer unwanted general differences in batch distributions.


**Note**: Use the automated rating for guidance only and in addition judge manually (e.g. by visual inspection of the totals overview) whether total concentration fits behave well enough in comparison to area or intensity (i.e. unnormalized and uncalibrated data). The rating remains rather subjective for now, as it will require a great number of experiments and extensive validation to assess which model coefficients and p-values are acceptable to support reliable analysis.


###### 4.2.1.2.1 Reference QCs


###### 4.2.1.2.1.1 Single batches

**Error:**: No p-value available for total Concentration [ï¿½M] slope of samples of type QC Level 2 in Batch **Batch1**! Too few samples available?

###### 4.2.1.2.1.2 Auxiliary Figures

###### 4.2.1.2.2 Biological samples


###### 4.2.1.2.2.1 Single batches

No batch with undesirable Concentration [ï¿½M] slope for samples of type Sample.

###### 4.2.1.2.2.2 Auxiliary Figures

###### 4.2.1.3 Missing values overview

##### 4.2.2 Well Plate Overview


**Warning:** Technical and biological variability can be separated, however the relation between the both is not fully conclusive.


Please see the details and the plot below for more information.

##### 4.3.2 Details

An integrative linear model is used to assess differences in technical and biological variability: `log(SD) ~ Group + log(Mean)/Group` with Group being a factor indicating QC or biological samples.

Model results are combined to identify four important main cases of differences in technical and biological variability:

1. `k >= 7`: A clear biological variability on top of the technical one.
2. `k = 2:6`: Inconclusive differences in biological and technical variability.
3. `k = 0:1`: A strong technical variability which hides the biological one.
4. `k < 0`: Something clearly wrong (when variability in QC decreases with increasing amount of metabolite concentration).

with final score k being computed to allow the classification of the four main cases as follows:

```
k = 0
k = k + (r_squared > 0.8)
k = k + 2*(delta_intercept > 0 & delta_intercept_pvalue < 0.05)
k = k + 4*(delta_slope > 0)
k = k - 8*(qc_slope < 0)
```

The results are as follows:

- Linear regression model results:
  - R squared = 0.9190357
  - QC slope = 1.0078533
  - Delta slope (Sample - QC) = -0.0874981
  - Delta slope p-value = 0.0872054
  - Delta intercept (Sample - QC) = 1.4199896
  - Delta intercept p-value = 1.483084710^{-14}
- Variability Score:
  - k = 3


##### 5.1.1 Pearson

###### 5.1.1.1 All sample types


###### 5.1.1.1.1 Type-colored labels

###### 5.1.1.2 QC samples


###### 5.1.1.2.1 Type-colored labels

###### 5.1.1.3 Biological samples


###### 5.1.1.3.1 Sex-colored labels

##### 5.1.2 Euclidean

###### 5.1.2.1 All sample types


###### 5.1.2.1.1 Type-colored labels

###### 5.1.2.2 QC samples


###### 5.1.2.2.1 Type-colored labels

###### 5.1.2.3 Study samples


###### 5.1.2.3.1 Sex-colored labels

#### 5.2 Missing value imputation

Median imputation is applied for the remaining visualizations on biological, reference and pooled QC samples (separately), as the methods are not suited to handle missing values. Samples which still contain missing values after imputation are completely removed.

#### 5.3 Sample correlation

Scatter plot matrices illustrate the correlation between samples based on compound concentration.

In the case of high sample numbers (> 9), several matrices will be generated based on sample groups of size up to 9 with one overlapping sample between each group (i.e. the last sample in a matrix is the first sample in the next matrix).

This analysis is based on the imputed dataset 4.

##### 5.3.1 Reference QCs


#### 5.3.1.1 1

##### 5.3.2 All samples


#### 5.3.2.1 1

#### 5.3.2.2 2

#### 5.4 PCA

Principal component analysis (PCA) is performed to emphasize compound concentrations-based sample type, batch and potential study group relationships.

#### 6.3 Parameters


- **additional\_normalization**: None
- **pqn\_reference\_type**: Sample
- **sample\_filter**:
- **metadata\_extraction**:
- **zero2na**: *TRUE*
- **ignore\_errors**: *TRUE*
- **data\_files**:

  - **Batch1**: /data/mkuhring/workspace/metaquac/inst/extdata/biocrates\_q500\_test\_01/Batch1\_LC.txt
- **kit**: Biocrates MxP Quant 500 Kit
- **measurement\_type**: LC
- **title**: Biocrates QC - Q500 - LC
- **author**: mkuhring
- **pool\_indicator**: Sex
- **profiling\_variables**: Sex
- **study\_variables**:

  - Sex
- **replicate\_variables**: Sex
- **preproc\_keep\_status**: Valid
- **filter\_compound\_qc\_max\_mv\_ratio**: *0.3*
- **filter\_compound\_qc\_max\_rsd**: *15*
- **filter\_compound\_bs\_max\_mv\_ratio**: *0.3*
- **filter\_compound\_bs\_min\_rsd**: *15*
- **filter\_sample\_max\_mv\_ratio**: *0.2*
- **data\_tables**: all
