## Supplementary material for "Concepts and Software Package for Efficient Quality Control in Targeted Metabolomics Studies – MeTaQuaC": QC Q500: qc_q500_2_fia.nb.html

Biocrates QC - Q500 - FIA


Code 

- Show All Code
- Hide All Code
- Download Rmd

### Biocrates QC - Q500 - FIA

###### *Mathias Kuhring*

###### *January 08, 2020*


This report was created with MeTaQuaC v0.1.1.

The data as imported, restructured and preprocessed in this report is available for export in the section Preprocessing (via the full data tables provided for each preprocessing step).

Before reading and interpreting this QC report, please make sure to be familiar with the Biocrates kit used, i.e. familiarize your self with the compounds, sample types, status values, terminology, analytical specification, etc.  
Please refer to Biocrates’ manuals and documents provided with the kit used.

### 1 Data Preparation


#### 1.1 Import

Import of **Biocrates MxP Quant 500 Kit** data of measurement type **FIA**.

Data files to import:


- **Batch1**: /data/mkuhring/workspace/metaquac/inst/extdata/biocrates\_q500\_test\_01/Batch1\_FIA.txt


Importing batch `Batch1`…

#### 1.2 Restructuring

The imported data was restructured as follows:

- Convert to **long table** (or molten dataset). Each row includes only one measurement, i.e. the data of one compound in one sample and all corresponding information.
- Add **Sample Identification “Blank”** for blank samples, as they are empty when exported from MetIDQ.
- Add **Sample Type “Pooled QC”** for pooled QC samples. Pooled QC samples are identified by any occurrence of the term “pool” (case-insensitive) in the column Sex (parameter `pool_indicator`). In contrast, **“Reference QC”** will refer to Biocrates’ QC Level 2 samples.
- Add **Sequence Position**, i.e. sample acquisition order. This position is calculated from MetIDQ’s Well Position, as this one depicts a position by row-major order which does not reflect the actual acquisition order.
- Add **Well Coordinates**, indicating the two dimensional position on the well plate. This position is calculated from MetIDQ’s Well Position and indicates rows by letters and columns by numbers.
- Add a **unique Sample Name**, which is a combination of Sample Identifier, Sequence Position and potentially Batch (if available). These names assure unambiguous identification and analysis of measurements within or between several batches (e.g. in particular technical samples such as calibration standard or reference QC samples share sample identifications).
- Remove **nonessential columns**, as some of them currently interfere with proper data merging or processing later. This includes (for now, to be optimized/reduced): Plate Bar Code, Sample Bar Code, Submission Name, Material, Plate Production No., Plate Note, Run Number, Injection Number, Measurement Time, Sample Description, Collection Date, Org. Info and OP.
- Unify **missing values**, i.e. “” (empty field), “0” (zero) and “NA” (not available) are converted to “NA”.
- **Factorize Compounds** with level order based on compound class and compound name.

### 2 Overview


#### 2.1 Samples and Compounds


##### 2.1.1 Samples


**Number of samples in total:** 17

**List of all samples:**


**Number of samples per batch:**

##### 2.1.2 Compounds


**Number of compounds in total:** 524

**List of all compounds:**


**Number of compounds per batch:**

Based on the parameter `preproc_keep_status`, measurements with the following statuses will be retained: **Valid**

Missing **Concentration** values before Biocrates status preprocessing: **2510** of **8908** (28%)

Missing **Concentration** values after Biocrates status preprocessing: **3916** of **8908** (44%)

#### 3.2 Step **2a**

2. **Remove unreliable **compounds**** (QC-based filters)
   1. Based on missing values ratio in Reference QCs (**>=30%**)

To alter the threshold use parameter `filter_compound_qc_max_mv_ratio`.

Number of compounds before: **524** Number of compounds left: **398**

#### 3.3 Step **2b**

2. **Remove unreliable **compounds**** (QC-based filters)
   2. Based on %RSD of Reference QCs (**>=15%**)

To alter the threshold use parameter `filter_compound_qc_max_rsd`.

Number of compounds before: **398** Number of compounds left: **318**

#### 3.4 Step **3a**

3. **Remove underrepresented **compounds**** (biological filters)
   1. Based on only missing values in biological samples (**=100%**)

Number of compounds before: **318** Number of compounds left: **297**

#### 3.5 Step **3b**

3. **Remove underrepresented **compounds**** (biological filters)
   2. Based on missing values ratio in biological samples (**>=30%**)

To alter the threshold use parameter `filter_compound_bs_max_mv_ratio`.

Number of compounds before: **297** Number of compounds left: **260**

#### 3.6 Step **3c**

3. **Remove underrepresented **compounds**** (biological filters)
   3. Based on %RSD in biological samples (**<15%**)

To alter the threshold use parameter `filter_compound_bs_min_rsd`.

Number of compounds before: **260** Number of compounds left: **213**

#### 3.7 Step **4**

4. **Remove underrepresented **biological samples**** (biological filters)
   1. Based on missing values ratio of compounds (**>=20%**)


**Note**: Expect inferior results for FIA data due to one-point calibration (in contrast to 7-point calibration in LC data).


###### 4.2.1.2.1 Reference QCs


###### 4.2.1.2.1.1 Single batches

**Error:**: No p-value available for total Concentration [ï¿½M] slope of samples of type QC Level 2 in Batch **Batch1**! Too few samples available?

The results are as follows:

- Linear regression model results:
  - R squared = 0.8745916
  - QC slope = 0.9788791
  - Delta slope (Sample - QC) = 0.0919488
  - Delta slope p-value = 0.0013547
  - Delta intercept (Sample - QC) = 1.2692933
  - Delta intercept p-value = 1.981037510^{-83}
- Variability Score:
  - k = 7

#### 6.3 Parameters


- **additional\_normalization**: None
- **pqn\_reference\_type**: Sample
- **sample\_filter**:
- **metadata\_extraction**:
- **zero2na**: *TRUE*
- **ignore\_errors**: *TRUE*
- **data\_files**:

  - **Batch1**: /data/mkuhring/workspace/metaquac/inst/extdata/biocrates\_q500\_test\_01/Batch1\_FIA.txt
- **kit**: Biocrates MxP Quant 500 Kit
- **measurement\_type**: FIA
- **title**: Biocrates QC - Q500 - FIA
- **author**: Mathias Kuhring
- **pool\_indicator**: Sex
- **profiling\_variables**: Sex
- **study\_variables**:

  - Sex
- **replicate\_variables**: Sex
- **preproc\_keep\_status**: Valid
- **filter\_compound\_qc\_max\_mv\_ratio**: *0.3*
- **filter\_compound\_qc\_max\_rsd**: *15*
- **filter\_compound\_bs\_max\_mv\_ratio**: *0.3*
- **filter\_compound\_bs\_min\_rsd**: *15*
- **filter\_sample\_max\_mv\_ratio**: *0.2*
- **data\_tables**: all
